# Biological influence of extracts of cryopreserved fragments of piglets’ heart and skin

**DOI:** 10.1101/184622

**Authors:** Liliia A. Rohoza, Iryna G. Bespalova, Mykola O. Chizh, Sergiy Ye. Halchenko, Boris P. Sandomirsky

## Abstract

One of the new directions on which the searches used to find the methods to effectively correct the regeneration in case of different pathologies is the application of biologically active peptides and their mixtures. In the work there was investigated the biological influence of extracts of cryopreserved fragments of skin and heart of newborn piglets with the cold wound of skin and myocardial ischemia in rats respectively. For investigations the extracts were obtained from cryopreserved fragments of newborn piglets’ skin and heart. Cold wound of skin was modelled in rats by 10 mm copper applicator cooled in liquid nitrogen down to -196°C; the areas of wounds were determined by planimetric method, the white blood cells’ counts were analysed. In rats with myocardial ischemia there were studied the electrocardiograms, heart rate variability and proliferative activity of heart cells. The injection of extracts of cryopreserved fragments of skin to the animal’s abdominal cavity accelerates the healing of cold wound of skin and normalizes the response of immune system to an injury. After the injection during 2 months to the animals with myocardial ischemia with extracts of cryopreserved fragments of heart the normalization of electrophysiological indices of heart activity was observed that testified about the improved blood supply to a heart muscle. Being injected to healthy animals and those with myocardial ischemia the extracts of cryopreserved fragments of heart resulted in an increase in proliferative activity of heart cells. The studied extracts have a high biological effect and can be applied when designing the drugs for regenerative medicine.

**Summary statement:** The extracts of cryopreserved fragments of piglets’ heart or skin were shown as stimulating reparative regeneration of heart tissues in myocardial ischemia of rats and skin in a cold wound, respectively.

## INTRODUCTION

Numerous investigations have convincingly proven that the main systems (nervous, endocrine, immune) which are responsible for maintaining the homeostasis and adequate response to various effects, have a common mechanism of chemical regulation. Underlying this mechanism is the production and secretion of a number of substances having a peptidic nature. These molecules have generally as regulatory peptides. Big attention in modern biology and medicine is paid also to the study of involvement of peptidic nature’s substances in regulation of homeostasis of some cell populations and to their role as the signal molecules, providing the communicative relationships in the norm and under pathological states (Rehfeld and Bundgaard eds, 2010). Physiologically active peptides were isolated virtually from all the inner organs, they have the ability of regulation of functional and proliferative activity of cells in the tissues, which are the initial material for their obtaining. Therefore the studying of their biological activity is deemed actual and important (Ivanov, 2010; Shpakov, 2013).

The peptidic bioregulators have been suggested to be a tissue specific. So, every complex of peptides can to specifically affect a physiological state of the tissues from which it has been obtained via the nonspecific influence to a body as a whole (Khavinson et al., 2014).

It was shown that the application of cryobiological techniques when obtaining the extracts from organ fragments of pigs and piglets allows to increase the yield and biological activity of peptides which containing in them (Rohoza et al., 2014a). There were matched the data obtained by MALDI-ToF method about the molecular masses of peptides in the extracts of cryopreserved fragments of piglets’ skin and heart with the known peptides from the Protein Knowledgebase (UniProtKB) (Rohoza et al. 2014b).

The research aim was to determine the biological influence of extracts of cryopreserved fragments of newborn piglets’ skin and heart in the cold wounds of skin (CWS) and myocardial ischemia (MI) in the rats respectively.

## MATERIALS AND METHODS

### Bioethics

The experiments were carried-out according to the “General principles of the experiments in animals”, approved by the 4rd National Congress in Bioethics (2010, Kiev, Ukraine) and coordinated with the statements of “European convention on the protection of vertebrate animals used for experimental and other purposes” (Strasbourg, 1985) as well as agreed by the Committee of bioethics of the Institute for Problems of Cryobiology an Cryomedicine of the National Academy of Sciences of Ukraine.

### Obtaining the extracts of cryopreserved organ fragments

Extracts of cryopreserved organ fragments were obtained from the skin and heart of newborn piglets. Hearts and skin were fragmented into 2-5 mg pieces and washed thrice with physiological solution (0.9% NaCl, pH 7.4) in ratio 1:10. To the fragments there was added dropwise the cryoprotective solution of 20% PEO - 1.500 in ratio 1:1, pre-packed into 20 ml plastic vials and frozen at 1 °C /min cooling down to -70°C with the following transfer into liquid nitrogen. Material was thawed on the water bath at 37-40 °C and cryoprotectant was washed out with physiological solution. For the extracts obtaining the fragments were incubated for 60 min in physiological solution. To remove thermolabile proteins the supernatant was warmed in water bath for 15 min and paper filtered. Concentration of peptides was measured spectrophotometrically at wave-length of 280 nm and 260 nm (Schmid, 2001).

### Experimental models

The tissue-specific influence of extracts of cryopreserved organ fragments was studied in outbred white male rats. Influence of the extract of cryopreserved fragments of newborn piglets’ skin (PiSE) investigated in the model of cold damage of rats’ skin, and influence of the extract of cryopreserved fragments of newborn piglets’ heart (PiHE) was studied in the rats with spontaneously myocardial ischemia (MI). Extracts were injected to rats into abdominal cavity once per day during the whole experiment. The dose of peptide was 500 μg per 1 kg of animal’s weight.

**Part 1.** Cold injury modelling. Cold wounds of skin (CWS) were modeled in rats of 180-210 g by 10 mm copper applicator cooled in liquid nitrogen down to - 196°C, the exposure was twice by 30 seconds. The animals were divided into 2 equal groups with 8 rats each: animals with the CWS which were injected with the physiological solution (control), and those with wound and injections with PiSE. The wound area was determined by digital images using Bio Vision software. To the 3d, 7th, 14th and 21^st^ days of experiment leukocyte counts on smears stained with azure II - eosin (Romanowsky-Giemsa) were analyzed. The index of blood leukocytes shift (IBLS) was counted as the ratio of the amount of eosinophils and neutrophils to the one of monocytes and lymphocytes.

**Part 2.** Animals with spontaneously arisen myocardial ischemia. Mature rats with MI were selected according to the analysis of ECG indices against the background of spontaneously arisen pathology under vivarium conditions. Age of the animals before the experiment was from 14 to 18 months. Totally there were examined 327 rats among those 18 rats with ischemia of the heart muscle were selected. For ECG recording the hardware-software complex “Poly-Spectrum” (Neurosoft, Russia) was used. The selection criterion was the elevation of ST-segment and increasing of T-wave amplitude, decreasing of R-wave amplitude in I and aVL leads. For estimation of rats’ organism regulatory systems according to ECG the analysis of the heart rate variability (HRV) was used. The group of norm (intact) was consisted of 18 rats without revealed pathological changes. The experimental group with ischemia of the heart muscle (10 rats) was injected with PiHE during 2 months. The control rats with ischemia of the heart muscle (7 animals) were injected with physiological solution in amount of 0.5 ml per 100 g of weight. Rats’ hearts for immune histological investigations were fixed in 10 % solution of the buffered neutral formalin (Shandon Fixx, USA) during 24 hrs. After dehydration the tissues were imbedded into paraffin with polymer additives (Richard-Allan Scientific, USA). Slices with 5mm thickness were made from the paraffin blocks. After blocking the nonspecific linking of proteins via the protein block (Diagnostic Biosystem, USA), primary antibody Ki-67 were applied (clone MIB-1, Code IR626, TRU FLEX, DAKO, Denmark). Primary antibody was visualized with the detection system DAKO EnVision FLEX+ (DAKO, Denmark). For visualization of histological structure the processed immunohistochemical samples of the heart were additionally stained with hematoxylin Mayer (DAKO, Denmark). To the sections which were used as a negative control instead the primary antibodies a buffer for antibodies dilution was used. The total number of Ki-67 positive cells were accounted 5 fields of vision (objective × 40, ocular × 10).

### Statistical analysis

The results were processed by the nonparametric method MANOVA with software SPSS Statistics 17.0. Data were expressed as mean ± the standard error of the mean.

## RESULTS

### Part 1. Healing of cold wounds, planimetry

Performed investigations for determining the healing rate of CWS in rats which were injected with PiSE and without injections shows that to the 3d day of experiment no statistically significant differences in the wound area in the control and experimental group were observed (Table 1). To the 7th day the wound area in the animals with PiSE injections was in 1.5 times less than in the control animals. As well to the 14th day it was also 3.8 times less than in the control. Thus beginning from the 7th day the wound area in the animals which were injected with PiSE was statistically significantly less than in the control rats. Thereby the studied extracts normalize the healing of cold wounds in the experiment.

**Table 1.** Cold wounds’ area (cm^2^) depending on experimental conditions.

| Time of observation, day | Experimental conditions |  |
| --- | --- | --- |
|  | Control<br>N=8 | Injections of PiSE<br>N=8 |
| 3 | 4.5±0.4 | 3.7±0.4 |
| 7 | 3.4±0.3 | 2.3±0.2* |
| 14 | 2.3±0.3 | 0.6±0.1* |
| 21 | 1.5±0.1 | Healing |

Note. * - differences are statistically significant compared to the control, p < 0.05

### Index of blood leukocytes shift (IBLS)

The level of the IBLS increasing under pathological states shows about activity of inflammation and disorder in immunological reactivity. To the 3d day after the CWS modeling the IBLS was considerably higher the norm in both groups of animals (Table 2). The most expressed differences of IBLS were to the 7th day of experiment. In the control group rats there was observed its further increasing from 1.01 up to 1.49, in the animals which were injected with the extract there was the IBLS decrease in contrast to the 3rd day. Following the PiSE injections to animals with the cold injury the IBLS value made 46% versus the control level. On the 14th day this index been decreased in the all groups of animals. To the 21st day in the animals injected with PiSE this index returned to the norm. This may testify that the injections of extracts reduces the inflammation manifestation and normalizes an immune response.

**Table 2.** Blood leucocytes shift depending on experimental conditions and observation time.

| Experimental conditions | Observation time, days |  |  |  |
| --- | --- | --- | --- | --- |
|  | 3 | 7 | 14 | 3 |
| Norm | 0.21 |  |  |  |
| Control | 1.01 | 1.49 | 0.70 | 0.29 |
| Cold wound + PiSE | 0.89 | 0.69 | 0.41 | 0.23 |

### Part 2. Analysis of heart rate variability

In the rats with MI injected with physiological solution during 2 months the HR, SDNN and CV practically do not change (Table 3). Nevertheless there was observed decreasing of the LF/HF ratio from 0.4 down to 0.3. The minimal and maximal recorded duration of cardiointervals in rats was to beginning of the experiment 100 and 493 ms and at the end of experiment it was 113 and 472 ms, respectively. In the group of animals with MI after PiHE injections during 2 months a recovery of R wave amplitude was recorded. Elevation of ST segment has been changed by appearance of a domed T wave that testifies about normalization of blood supply to the heart muscle. In this group of animals among all of the studied indices of HRV only the HR was not statistically and significantly differ from the HR of healthy animals; average HR increased from 381 up to 498 bits per min. To the experiment beginning the minimal recorded cardiointerval was 127 ms and at the end it was 95 ms. The maximal cardiointerval was 497 and 148 ms respectively. In the norm these indices were 108 and 144 milliseconds. The clearly expressed normalization of the rest of the indices was observed. Thus, if to the experiment beginning the SDNN value was higher the norm in 5.6 times so at the end it was only 1.5 times higher, and its value decreased in 6.7 times. The initial coefficient of variation exceeds the norm in 12.4 times and after the PiHE injections during 2 months this rise made just 1.8 times. The LF/HF ratio increased from 0.4 up to 2.8 versus the norm rate of 5.6. After 2 months of injections with the extracts the SDNN, LF and HF values did not statistically and significantly differ from these indices in the norm. Thus the analysis of HRV revealed that in the animals with the heart muscle ischemia there is observed significant decrease in the LF/HF ratio and, hereby, also the preference of parasymphatetic influences upon the heart. PiHE injection to animals with such a pathology contributes to normalizing all the examined in this research indices.

**Table 3.** Indices of the heart rate variability in rats with myocardial ischemia.

| Indices | Animal groups |  |  |  |  |
| --- | --- | --- | --- | --- | --- |
|  | Injections with saline<br>N=6 |  | Injections with PiHE<br>N=6 |  | Norm<br>N=24 |
|  | Initial<br>indices | After 2<br>months | Initial<br>indices | After 2<br>months |  |
| HR,<br>beats per<br>min | 394±31 | 407±35 | 381±29 | 498±37<sup>1,2</sup> | 523±45 |
| Minimum<br>duration of<br>RR<br>interval, ms | 100 | 113 | 127 | 95 | 108 |
| Maximum<br>duration of<br>RR<br>interval, ms | 493 | 472 | 479 | 148 | 141 |
| SDNN | 53.4±4.4 | 50.2±4.6 | 50.8±5.1 | 13.4±1.1<sup>1,2</sup> | 9.9±0.6 |
| CV, % | 24.1±2.0 | 21.2±1.9 | 26.1±2.6 | 3.8±0.2<sup>1</sup> | 2.6±0.2 |
| LF, % | 28.6±2.5 | 25.5±2.1 | 27.8±2.9 | 74.8±7.1<sup>1,2</sup> | 79.1±6.4 |
| HF, % | 72.7±5.4 | 75.9±6.6 | 73.3±7.9 | 26.1±2.2<sup>1,2</sup> | 21.2±1.8 |

Note. 1 - differences are statistically significant if compared to the initial indices, p < 0.05;

2 - differences are not statistically significant if compared with norm, p > 0.05

### Proliferative activity

In intact animals with no detected pathology of the heart the number of Ki-67 positive cells was 7.9% of the total amount and in the animals with MI - 12.0% (Table 4). Injections with physiological solution both to intact animals and those with heart pathologies did not influence the number of these cells. In healthy animals injected with PiHE the number of Ki-67 positive cells in myocardium after 2 months of the injections was 6.1%. In the rats with MI at this term of observation the number of these cells was (24.6%).

**Table 4.**
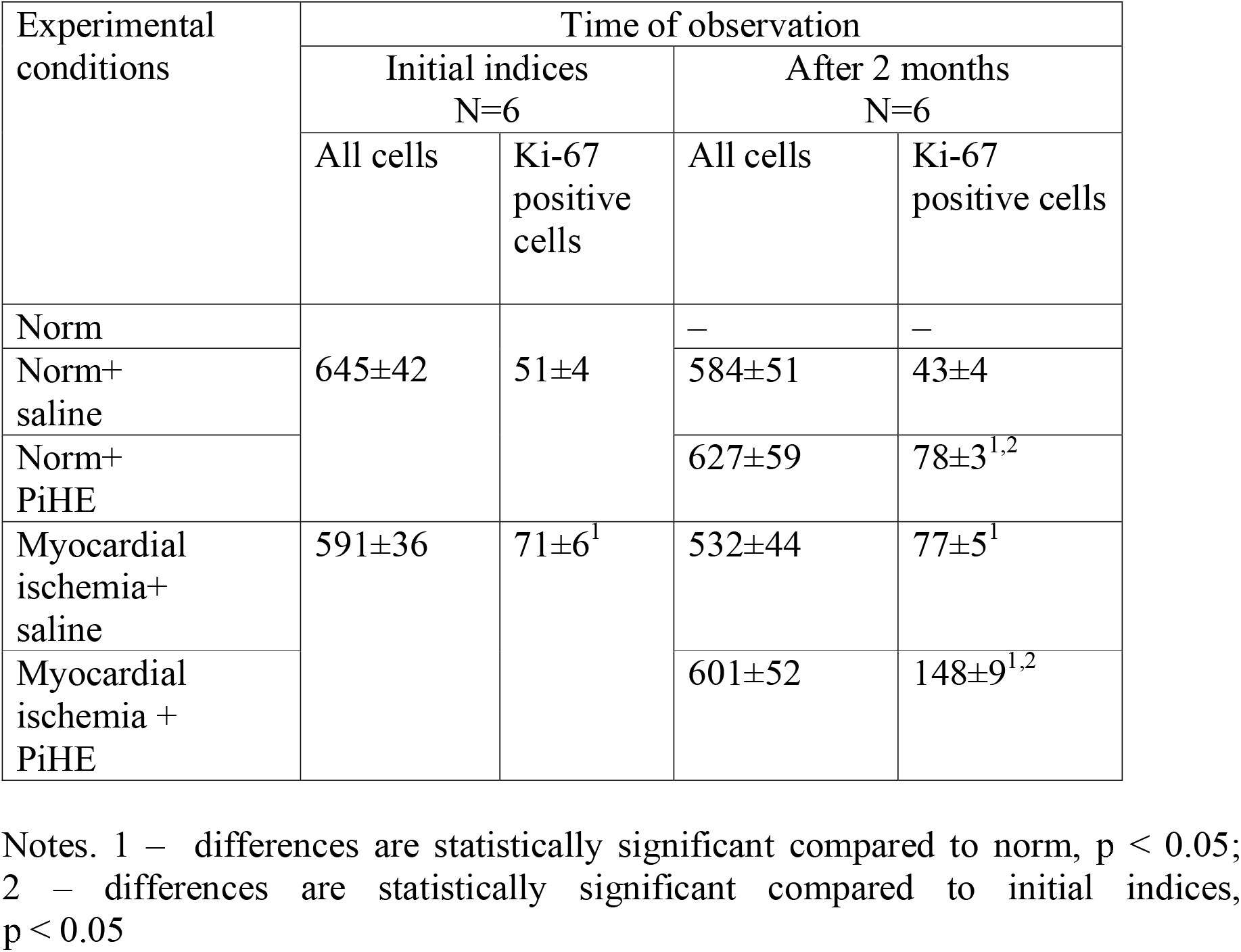
Total quantity and quantity of Ki-67 positive cells in myocardium of rats

1 - differences are statistically significant compared to norm, p < 0.05;

2 - differences are statistically significant compared to initial indices, p < 0.05

Thereby, the injection with PiHE increases proliferative activity of cells in the myocardium in the same way it stimulates the reparative regeneration of heart which is physiologically inherent to heart muscle.

Thereby the PiSE injection into animal’s abdominal cavity accelerates the healing of CWS and normalizes an immune response in trauma. After PiHE injections during 2 months to the animals with MI there were observed an increase in HR, recovery of R wave amplitude in ECG, elevation of ST segment was changed by appearance of a domed T wave testifying to normalization of blood supply to the heart muscle. The coefficient of variation of heart rhythm decreased in 6.7 times, and the ratio of LF/HF increased from 0.4 up to 2.8. Herewith there was observed an increase in the index of the spectrum power of neurohumoral regulation for all the frequency bands up to the normal level and restoration in the balance of contributions of sympathetic and parasympathetic divisions of autonomic nervous system into the heart rate regulation.

The PiHE injection to healthy animals and those with MI led to increasing the proliferative activity of myocardium cells.

## DISCUSSION

Studying the mechanisms of biological influence of the tissue-specific peptides is an actual task of current molecular biology, physiology and medicine. The main functions of these peptides are the control of proliferation, differentiation and elimination of cells from a relevant tissue. It is specific, that their effect herewith is presented not by the individual components, but by large number of peptides with variously directed activities. The methods traditionally applied in combined treatment of wounds (antibiotics, sorbents and others) and various physical influences to the wound not always allow a sufficient normalization of healing. For this reason recently there have been re-considered the approaches to wound treatment (Sood et al., 2014; Murphy and Evans, 2012; Sarabahi, 2012). One of the novel directions when searching the methods for effective correction of reparation is the use of different means for regulation of wound healing. For wounds treatment there are used biologically active substances, such as cytokines, hormones, neuropeptides, regulatory peptides and synthetic peptides (Evans et al., 2013; Mu et al., 2014; Borena et al., 2015; Tadokoro et al., 2015; You and Han, 2014).

Application of the abovementioned active agents enables to balance the inflammation and regeneration whereat the wound healing occurs and also to control the course and change of the different stages of these processes. However, when using the physiologically active medications of mono-component peptides the impact only on a certain link of the regeneration process occurs. Instead, the poly-component mixtures containing a wide range of regulatory peptides may influence several key links of this process simultaneously (or sequentially). Especially this is important under conditions of an infected wound with strong bacterial contamination, pronounced disorders of metabolism, microcirculation and lymphodynamic under which there is marked a discoordination in the system of autoregulation of wound process. It is noted that thereat the spectra of cytokines secretion are changing. There are dominated the inhibitory rather than inductive factors, an insufficient inflow of systemic regulatory peptides to wound is observed (Schreml et al., 2016; Eming et. al., 2007). Such a damage promotes a prolonged flowing for all the wound phases, lasting healing of wounds and formation of keloid scars. Namely the necessity to eliminate the indicated disorders in the system of peptidic regulation of wound healing enables to pathogenetically substantiate the application in a combined treatment of wounds the medications of immunobiological effect. In recent decades an interest to applying the regulatory peptides in cardiology has been considerably increased too. In particular, such a problem as stimulation of reparative regeneration of myocardium under the ischemic heart disease (IHD) is an acute one and constantly updated by the new approaches of its resolving. Currently there are extensively investigated the effectiveness and mechanisms of action involving the cell therapy, growth factors and various peptides as well (Segers and Lee, 2008; Williams and Hare, 2011; Yla-Herttuala et al., 2007; Hayek and Nemer, 2011). These studies require not only advanced experimental and clinical investigations, but also general conceptions as for the mechanisms affecting regeneration.

Traditional treatment and secondary preventive measures of IHD represent a single set of measures, including therapy with medication, in particular application of antianginal, and if necessary also antiarrhythmic drugs, cardiac glycosides etc. (Ahmed et al., 2008; Dobrev and Nattel, 2010; Shavelle, 2007; Dai and Ge, 2012). The necessity of long-term using of these drugs makes the doctors fairly uneasy about their application safety, because according to the pharmacological properties of drugs in patients may appear the side effects. Conducting an adequate antithrombotic therapy may provoke bleeding, particularly in patients with existing lesions of gastrointestinal and with the presence of other risk factors. Also might be increasing the activity of transaminases (Calderon et al., 2010). This once again proves a need of searching a new alternative and even more effective treatment methods of IHD which enable not only overcome the problem, but also avoid some of side effects.

Number of papers already investigated the effect how the mono-, as the polycomponent peptidic drugs under various pathological states of the heart in vivo and in vitro. Under effect of an exogenous peptide apelin-12 there were observed the functional recovery of post-ischemic heart and preservation of cell membranes, resulted from an aerobic metabolism improvement and anticoagulant protection of myocardium (Pisarenko et al., 2012). When using the peptide apelin-13 in the models of myocardial infarction and damages under ischemia/reperfusion of the heart in animals its cardioprotective effect was shown (Azizi et al., 2013). As well it has been shown that when injecting into a body the mixtures of peptides under hypercoagulation conditions there was observed a stimulation of functions of anticoagulant system and fibrinolytic activity (Grigorjeva and Pyapina, 2010) etc. These polycomponent mixtures are more perspective and have been intensively studied.

**List of symbols and abbreviations**
CV: coefficient of variation
CWS: cold wound of skin
ECG: electrocardiograms
HF: high frequency component of HRV
HR: heart rate
HRV: heart rate variability
IBLS: index of blood leukocytes shift
IHD: ischemic heart disease
LF: low frequency component of HRV
MALDI-ToF: time-of-flight matrix-assisted laser desorption / ionization MI myocardial ischemia PEO polyethylene oxide
PiHE: extracts of cryopreserved fragments of heart of newborn piglets
PiSE: extracts of cryopreserved fragments of skin of newborn piglets
SDNN: standard deviation of the average duration of RR-intervals (NN-intervals)

## Acknowledgements

The authors express their gratitude to the Head of Animal Hosue Facility at the Institute for Problems of Cryobiology and Cryomedicine of the NAS of Ukraine Lyudmyla Batsunova for her assistance in handling with animals. We are also thankful to Dr. Kateryna McDonald, School of Health, University of Central Lancashire, UK for productive discussion and her valuable help in preparing this manuscript.

## Competing interests

The authors have no conflicts of interest and financial obligations.

## Author contributions

B.S. - interpretation of the findings, supervision of the experiment design; S.H. - planning the experiments, deriving the extracts, analysis of the obtained results; M.Ch. - participation in the experiments on influence of the PiHE injections to the rats with MI on electrophysiological indices of heart activity; L.R. - investigation of the effect of PiHE injection on proliferative activity of rats’ heart cells and PiHE injections to the animals with MI on electrophysiological indices of heart activity; I.B. - providing the experiments on the influence of the PiSE injections on the cold wound healing.

## Funding

Funding was provided by the National Academy of Sciences of Ukraine on the topic registered as “Destructive and renewing processes in the tissues in vivo after effect of biologically active components» to be performed at the Department of Experimental Cryomedicine”.

